# Junctional β-Catenin Stabilization Links Wnt Signaling and Force Generation

**DOI:** 10.64898/2026.05.06.723156

**Authors:** Agimaa Otgonbaatar, Sujithra Shankar, Prameet Kaur, Prabhat Tawari, Nicholas S. Tolwinski

## Abstract

β-catenin plays two fundamental roles in animal tissues: it acts as a transcriptional effector of canonical Wnt signaling and as a core structural component of adherens junctions that mediate cell–cell adhesion. In canonical Wnt signaling, the post-transcriptional regulation of β-catenin abundance, primarily through regulated phosphorylation, ubiquitination, and proteasomal degradation, determines whether the pathway is “off” or “on.” Despite the central importance of β-catenin stabilization, *in vivo* measurements of β-catenin protein lifetime and stabilization dynamics during development remain limited. Here, we measure the stability of endogenous β-catenin *in vivo* using tandem fluorescent protein timers (tFPs; “Timers”) inserted as minimally disruptive cassettes within the endogenous locus. Timers allow simultaneous visualization of a newly synthesized, rapidly accumulating pool (fast-maturing GFP) and a long-lived, stabilized pool (slow-maturing RFP). Surprisingly, the strongest stabilization does not occur in canonical Wnt patterning stripes; instead, we observe marked stabilization of junctional β-catenin at the leading edge during dorsal closure, a force-generating morphogenetic process. This stabilization is not explained by canonical Wnt ligand input and seems to reflect a stability program linked to β-catenin’s adhesive function in adherens junctions. We suggest that a stable junctional pool of β-catenin is vital for dorsal closure mechanics and provide evidence that this stabilization is regulated by Dishevelled and JNK, thus connecting Wnt pathway components to mechanotransduction.

## INTRODUCTION

### β-catenin at the intersection of adhesion, signaling, and morphogenesis

The molecular basis of multicellularity is inseparable from the evolution of stable cell– cell adhesion. The discovery of cadherins as transmembrane adhesion molecules^1,2^ and the demonstration that their intracellular domains are required for stable adhesion^3^ established the framework for adherens junctions. Cadherins associate with intracellular catenins—α-catenin, β-catenin, and γ-catenin—forming a multi-protein complex that links cell–cell adhesion to the actin cytoskeleton.^4^ In this complex, β– and γ-catenin bind directly to the cadherin cytoplasmic tail,^5–7^ whereas α-catenin, which is homologous to actin-binding proteins, supports cytoskeletal coupling.

Historically, β-catenin was first recognized as a structural component in adherens junctions. This perspective changed considerably when β-catenin was identified as the vertebrate equivalent of the *Drosophila* segment polarity gene *armadillo,*^8–11^ a gene discovered through classic developmental patterning screens.^12,13^ Later, as β-catenin was confirmed as the main cytoplasmic-to-nuclear effector of canonical Wnt signaling, the field’s focus shifted to signal transduction: pathway components, post-translational modifications, transcriptional partners, and downstream gene targets. A lingering question remained: is β-catenin primarily a membrane or cytoplasmic protein with occasional nuclear signaling, or does its signaling activity arise from a separate, distinct pool?

### Dynamic adherens junctions and the Wnt–polarity–force interface

Adherens junctions are now understood to be dynamic and spatially regulated rather than purely structural. In *Drosophila* embryos, junctions are redistributed asymmetrically during convergent extension^14,15^ and are regulated by epithelial polarity^16,17^ and influenced by non-canonical Wnt polarity signaling^18,19^. Across systems, junction remodeling is tightly coupled to force-generating machinery and morphogenesis.^20–25^ This raises a key question for β-catenin biology: are β-catenin’s nuclear and junctional roles separate outputs, or do they converge through shared regulatory mechanisms during mechanically active tissue remodeling?

### Measuring β-catenin dynamics in vivo with engineered endogenous alleles

Several technical advances now enable direct in vivo analysis of protein dynamics at endogenous levels. Genome engineering allows modification of endogenous genes to insert intragenic cassettes encoding fluorescent proteins or functional modules.^26–30^ Optogenetics enables perturbation of protein behavior with light, enabling temporally controlled functional assays. In our earlier work, we used overexpression in normal and mutant backgrounds to probe signaling dynamics and timing.^31–33^ By combining genome engineering and optogenetic tools, we can now both observe and perturb pathway components in living embryos.^34,35^

A persistent challenge is that endogenously tagged β-catenin reporters often primarily reveal the abundant membrane-associated pool and are less effective at reporting the canonical signaling pool *in vitro*.^36,37^ To overcome this limitation, we inserted a tandem fluorescent protein timer into the endogenous *armadillo* gene. Timers combine a fast-maturing fluorescent protein (here, superfolder GFP) with a slow-maturing RFP, enabling inference of protein lifetime and stability from the GFP/RFP ratio^38–42^.

### Canonical Wnt regulation of β-catenin

In canonical Wnt signaling, β-catenin is the central relay from the membrane to the nucleus.^43^ In the absence of Wnt ligand, cytoplasmic β-catenin is phosphorylated by CK1 and GSK3 and targeted for ubiquitination by the destruction complex (Axin, APC, CK1, GSK3), keeping cytoplasmic β-catenin levels low.^44,45^ When Wnt binds Frizzled (Fz) and LRP5/6 (Arrow in *Drosophila*), Dishevelled (Dsh) is recruited to the membrane and nucleates assembly of the Wnt signalosome, recruiting Axin and kinases. This disables the destruction complex, allowing β-catenin to accumulate, enter the nucleus, and activate transcription through TCF.^46,47^

Importantly, β-catenin’s junctional and signaling roles are both regulated by phosphorylation at distinct sites. Destruction-complex phosphorylation targets β-catenin for degradation. Other kinases (e.g., JNK, Src) have been proposed to regulate junctional function by phosphorylating β-catenin and thereby modulating cadherin association.^48,49^

### Dorsal closure: a force-driven morphogenetic context to study junctional β-catenin stabilization

We focus on dorsal closure, a classic morphogenetic process in embryogenesis. After germ-band retraction, an epidermal gap covered by the amnioserosa occupies the dorsal side (Stage 13). During dorsal closure, lateral epidermal sheets converge and the amnioserosa contracts, closing the dorsal opening and restoring a continuous epidermis (Stage 17).^50–52^ The leading edge of the epidermis forms an actin cable coupled through adherens junctions. Signaling and force-generating processes must be coordinated, and Armadillo is required. Mutations in *arm* and *wg* show dorsal closure defects^53,54^, but the mechanistic basis remains unclear. Here, we ask why β-catenin/Arm is strongly stabilized at the leading edge during dorsal closure and what molecular mechanisms govern this stabilization.

## RESULTS

### Arm localizes to the leading edge and becomes highly stabilized during dorsal closure

Attempts to visualize endogenously tagged β-catenin primarily detect membrane– and cytoplasmic pools and do not robustly report the nuclear/signaling pool *in vitro*, including during embryogenesis, where expected Wnt stripes are subtle or absent with conventional exon traps.^36,37^ We therefore enhanced endogenous visualization by inserting a tandem fluorescent protein timer (tFP) into *arm*. The timer uses fast-folding superfolder GFP and slow-maturing RFP, so that GFP highlights newly synthesized or recently accumulated protein, while RFP indicates protein that has persisted long enough to mature—an operational readout of increased stability/longer lifetime.^38–42^

*In toto* imaging revealed broadly distributed Arm-GFP. However, the earliest clear appearance of a stabilized (RFP-positive) pool occurred specifically in the first row of epidermal cells bordering the dorsal opening—the leading edge—during dorsal closure. The combined GFP and RFP signals at the leading edge produced a bright yellow overlap, indicating that Arm is not only abundant but also strongly stabilized in these cells during dorsal closure (Figure 1).

**Figure 1.**
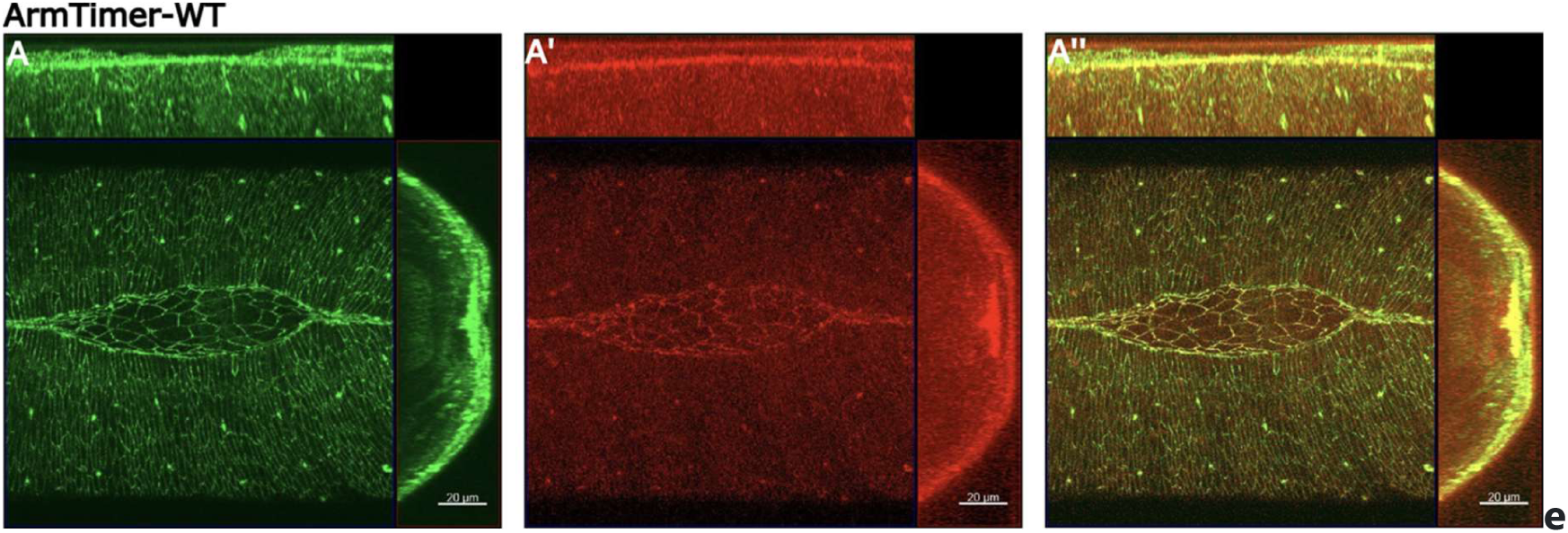
The arm stabilizes at the leading edge. (A) Green fluorescence indicates newly accumulating Arm, as only the GFP has had time to mature. Higher levels of Arm localize at the leading edge (the first row of epithelial cells of the dorsal gap). (B) Red fluorescence shows stable Arm protein with time for RFP to mature. (C) The leading edge exhibits bright yellow fluorescence, showing colocalization of mainly within dorsal closure.

### Arm activity is necessary for dorsal closure

Previous genetic studies reported dorsal closure defects in arm mutants.^54,55^ However, strong arm loss often disrupts epithelial cohesion earlier in development, complicating interpretation because the adhesive and signaling functions cannot be separated when the epithelium fails prematurely.^56,57^

To test Arm function with temporal control, we inserted an optogenetic cassette (Cryptochrome 2 and mCherry) into *arm* using MiMIC/RMCE. Blue-light exposure (488 nm) induces CRY2 oligomerization, which functionally inactivates Arm fused to CRY2.^31^ Arm-CRY2-mCherry localized normally to the leading edge in the absence of activation (Figure 2A), indicating that the insertion did not disrupt localization or baseline function. In contrast, blue-light activation caused a severe dorsal closure defect (Figure 2A’), demonstrating that Arm activity is required during dorsal closure. While this result is consistent with an adhesive role, it does not, on its own, distinguish adhesion from signaling.

**Figure 2.**
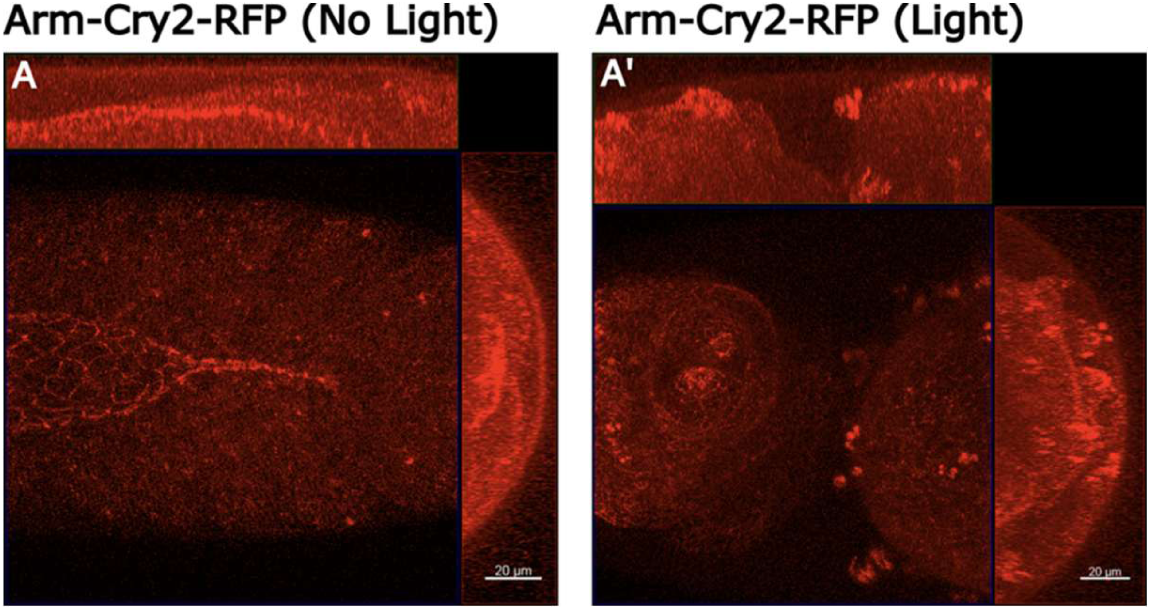
Arm is required for dorsal closure. (A) Arm-CRY2-mCh without blue-light activation localized at the leading edge of dorsal closure. The insertion of CRY2 and RFP did not perturb its localization or function. (A’) Light-induced oligomerization of Arm-CRY2-mCh through the application of blue light led to a disruption in dorsal closure.

### Upstream activation of canonical Wnt does not affect dorsal closure or the stabilized Arm pool

Embryos lacking Wnt signaling (e.g., wingless mutants) exhibit dorsal closure defects with an anterior break in zippering.^54^ We therefore tested whether increasing Wg ligand levels could enhance Arm stabilization at the leading edge and/or perturb dorsal closure.

In wild-type embryos, Wg is expressed in anterior–posterior stripes^58^ oriented perpendicular to the dorsal closure leading edge, where stable Arm accumulates (Figure 3A). Overexpression of Wg specifically in the amnioserosa (C381-Gal4) did not disrupt dorsal closure (Figure 3B). Uniform overexpression of Wg likewise did not alter Arm-RFP levels at the leading edge or cause a dorsal closure phenotype (Figure 3B’). These results indicate that the stabilized leading-edge Arm pool is not simply increased by ectopic upstream canonical Wnt ligand input.

**Figure 3.**
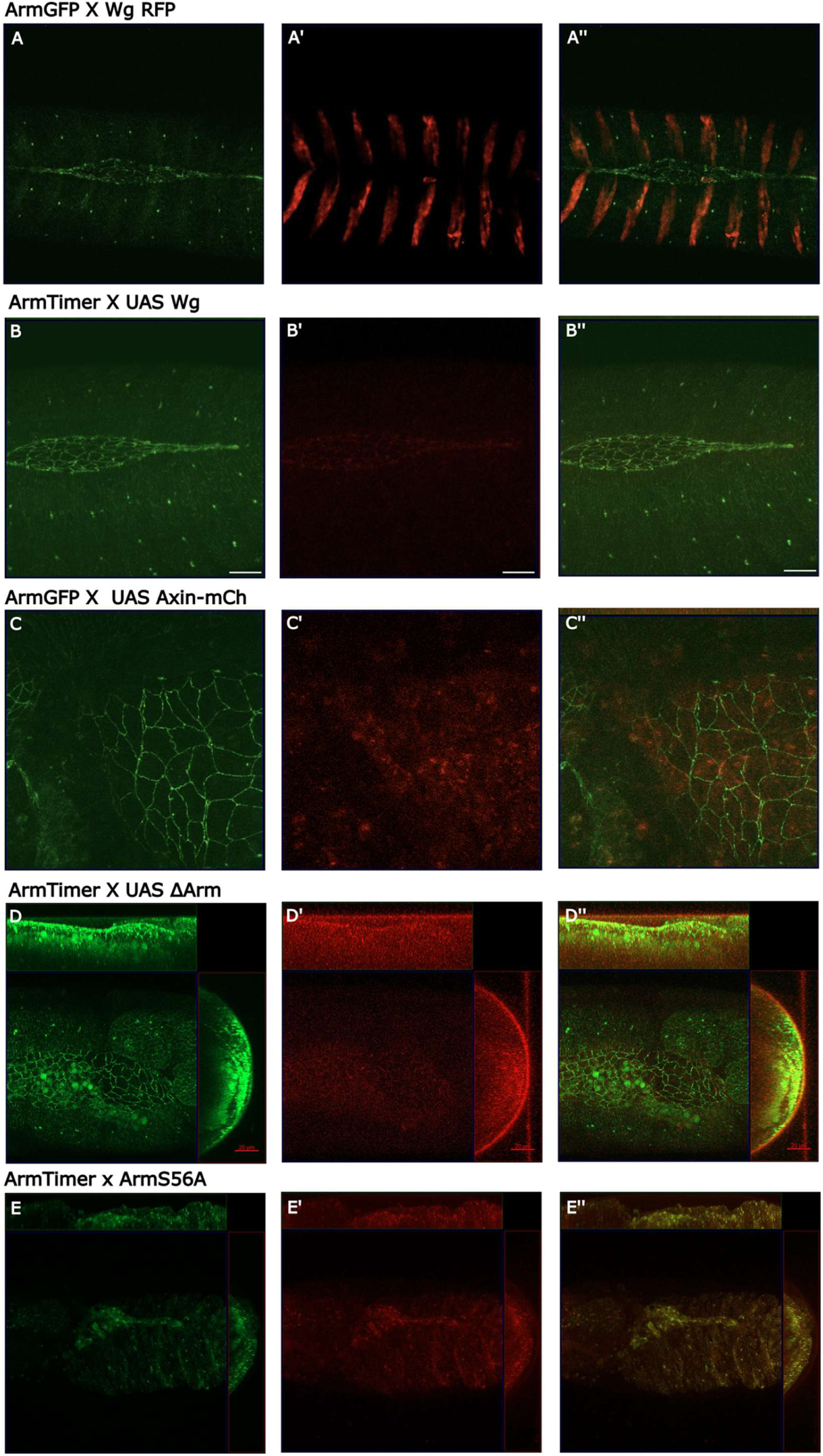
Canonical Wnt activation does not affect dorsal closure, while inactivation of Wnt results in defective dorsal closure. (A–A’’) Wingless (C381-Gal4 X UAS-Wg, UAS-RFP), the primary Wnt pathway ligand, is expressed perpendicular to the leading edge cells (Arm-GFP), and does not affect dorsal closure. (B–B’’) Wnt activation through uniform Wg overexpression (UAS Wg) does not affect dorsal closure. (C–C’’) Wnt signaling inactivation through Axin overexpression results in dorsal closure defect. (D–D’’) Expression of the gain-of-function allele of Arm, ΔArm resulted in a dorsal closure defect. (E–E’’) Gain of function allele ArmS56A also showed a dorsal closure defect.

### Downregulating canonical Wnt downstream, or forcing Arm stabilization, causes dorsal closure defects

Because Wg overexpression did not affect Arm stabilization or dorsal closure, we tested inhibition downstream at the level of Arm stability control. Axin is a rate-limiting scaffold of the β-catenin destruction complex.^44,59^ Overexpression of Axin prevents Arm stabilization and inhibits Wnt signaling. Axin overexpression produced a strong dorsal closure defect (Figure 3C), indicating that perturbing the stability machinery can disrupt dorsal closure.

We then tested stabilized versions of Arm, or those resistant to phosphorylation-dependent degradation. ΔArm lacks the N-terminus required for destruction-complex phosphorylation and degradation.^60^ ArmS56A is a gain-of-function allele that blocks phosphorylation by substituting the first serine with alanine in the degradation motif, thereby preventing normal degradation. Expression of either ΔArm or ArmS56A caused severe dorsal closure defects (Figure 3D,E). Because Wg overexpression did not cause a phenotype, these results are consistent with the possibility that dorsal closure is sensitive to direct perturbations of Arm stability/structure (bypassing normal regulation) rather than to ligand-mediated canonical activation. Additionally, ΔArm lacks the α-catenin-binding site, suggesting that impaired junctional coupling contributes to the phenotype.

### Adherens junction components co-localize at the leading edge, and α-catenin is required for dorsal closure

To assess whether leading-edge Arm stabilization reflects adherens junction reinforcement, we imaged E-cadherin during dorsal closure and observed strong colocalization with Arm at junctions (Figure 4A). We also visualized the actin cytoskeleton with LifeActin-mKO and observed the characteristic actin cable and distribution aligned with the leading edge, consistent with force generation and junctional coupling during zippering (Figure 4B).

**Figure 4.**
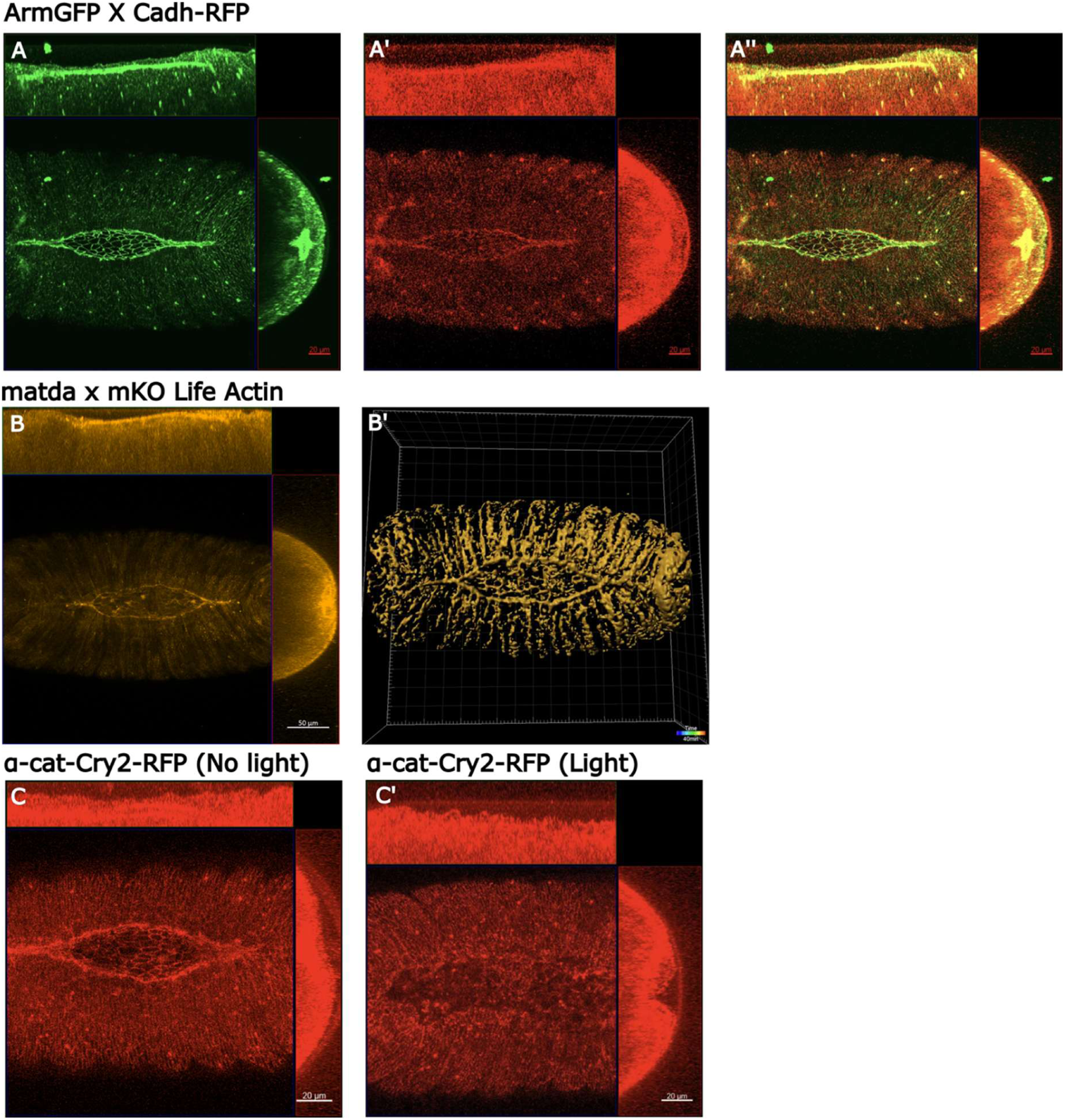
E-cadherin, actin and α-catenin co-localize with Arm at the leading edge, and α-catenin is necessary for dorsal closure. (A–A’’) E-cadherin (red) colocalizes with Arm (green) at junctions. (B, B’) Filamentous actin follows a similar pattern to adhesive junctions (B mKO tagged LifeActin, B’ projection using IMARIS software). (C) In the absence of blue light, α-catenin (red) localizes at the leading edge. (C, C’) Activation of CRY2 with blue light led to a loss of localization of α-catenin and a severe dorsal closure defect.

Because genetic disruption of adherens junction components often causes early defects that complicate interpretation during dorsal closure, we used an optogenetic approach targeting α-catenin. We inserted CRY2 and mCherry into α-catenin via MiMIC-RMCE, generating α-cat-CRY2. Without activation, α-catenin localized normally to the leading edge, and dorsal closure proceeded (Figure 4C). Blue-light activation reduced α-catenin activity and caused dorsal closure failure (Figure 4C’). Notably, the α-cat-CRY2 phenotype closely resembled Arm-CRY2, ΔArm, and ArmS56A phenotypes, particularly in the failure of zippering forces at the two closure edges. Together, these results support the conclusion that adherens junction function—specifically involving Arm and α-catenin—is essential for dorsal closure.

### Arm point mutants indicate that Arm–α-catenin interaction is required for dorsal closure

To dissect the mechanism underlying Arm function during dorsal closure, we engineered point-mutant *arm* alleles. We used MiMIC/RMCE to knock in mutated Arm alleles fused to the Timer construct into the first coding intron, followed by a stop codon to prevent translation of the endogenous gene. We validated this method with a hypomorphic Arm allele (ArmF1a), which has reduced activity but normal stability.^59^ Imaging ArmF1a-Timer revealed a mild dorsal closure defect in which the dorsal gap fails to fully close, supporting the effectiveness of this allele-replacement strategy (Figure 5B).

**Figure 5.**
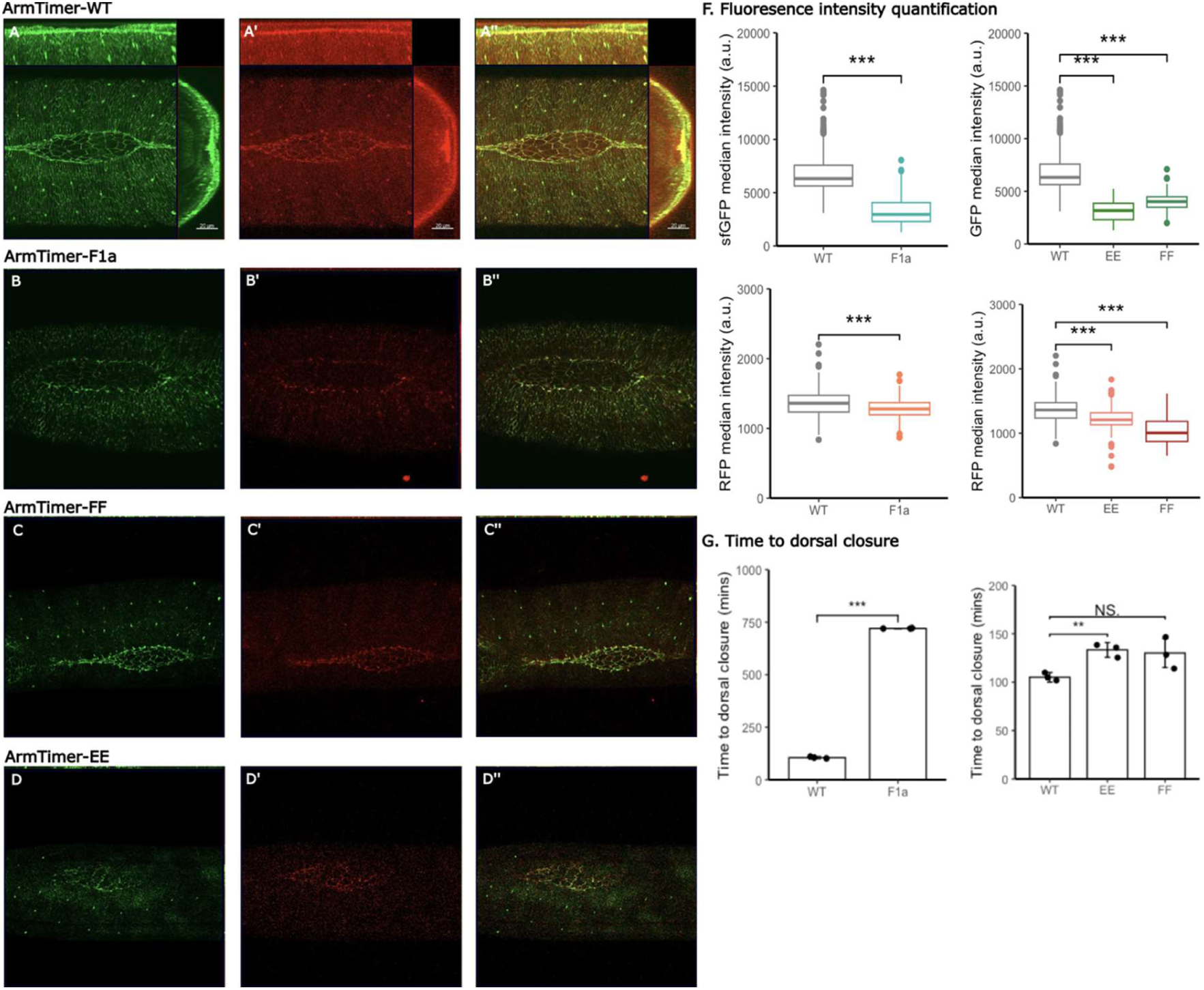
Functional dissection of point mutations in ArmTimer during dorsal closure. (A–A’’) Wild-type ArmTimer control shows high production and stabilization at the leading edge, as indicated by green (GFP) and red (RFP) fluorescence. (B–B’’) ArmTimer-F1a shows a mild dorsal closure defect and reduced fluorescence, validating the allele-replacement strategy. (C–C’’) ArmTimer-FF (Y150F, Y667F) shows no overt dorsal closure defect but reduced GFP and RFP intensities. (D–D’’) ArmTimer-EE (Y150E, Y667E) shows no overt defect but further reduced GFP and RFP intensities. (F) Quantification of GFP/RFP intensities comparing wild type, –EE, and –FF. (G) Dorsal closure duration is increased in ArmTimer-EE.

We next examined four phosphorylation sites: two tyrosines (Y150, Y667) implicated in regulating E-cadherin binding and junctional dynamics^49,61–65^, and two threonines (T111, T121) within the α-catenin-binding region that influence Arm–α-catenin binding.^16,66^ Each engineered allele includes the Timer tag to enable simultaneous visualization of “new” (GFP) and “stabilized” (RFP) pools.

For the tyrosine sites, we generated two variants:

- ArmTimer-FF, where Y150 and Y667 are replaced with phenylalanine, preventing phosphorylation at these sites (phospho-resistant).
- ArmTimer-EE, where Y150 and Y667 are replaced with glutamic acid, intended to mimic phosphorylation by introducing a negative charge (phosphomimetic).

Both ArmTimer-FF (Figure 5C) and ArmTimer-EE (Figure 5D) completed dorsal closure without gross morphological defects, yet both showed markedly reduced fluorescence intensity compared to wild-type ArmTimer. Quantification confirmed that both mutants exhibited significantly reduced GFP and RFP signals (Figure 5F). Notably, the pattern of reduction differed between the two mutants: ArmTimer-EE showed lower GFP intensity but comparable RFP intensity relative to wild type, whereas ArmTimer-FF exhibited reduced RFP intensity but similar GFP intensity compared to wild type. Interpreting the timer signals, this suggests that ArmTimer-EE may reduce overall production/accumulation (lower GFP) while preserving a stabilized pool (RFP), whereas ArmTimer-FF more strongly affects stabilization (reduced RFP) without strongly affecting the rapidly accumulating pool (GFP).

Although these tyrosine mutants did not cause overt closure failure, they did influence the duration of dorsal closure (Figure 5G). Specifically, ArmTimer-EE showed a significantly longer closure duration compared to wild type, while ArmTimer-FF did not significantly alter closure timing.

In contrast to the tyrosine mutants, altering the threonine sites within the α-catenin binding region produced a clear functional phenotype. ArmTimer-AA (T111A, T121A), which prevents phosphorylation at these threonines and reduces affinity for α-catenin, disrupted dorsal closure dynamics (Figure 6). The phenotype was characterized by defective and asymmetric zippering, resulting in an uneven dorsal gap and a substantially prolonged closure process compared to wild-type ArmTimer. Both ArmTimer-AA and ΔArm disrupt Arm–α-catenin interaction, but to different extents: ΔArm removes the binding region entirely, while ArmTimer-AA reduces binding affinity by modifying two regulatory sites. The graded phenotypes match this expectation: ΔArm causes failure of closure, while ArmTimer-AA produces slower and asymmetric zippering rather than complete failure. Together, these results map dorsal closure robustness onto the integrity of Arm–α-catenin coupling.

**Figure 6.**
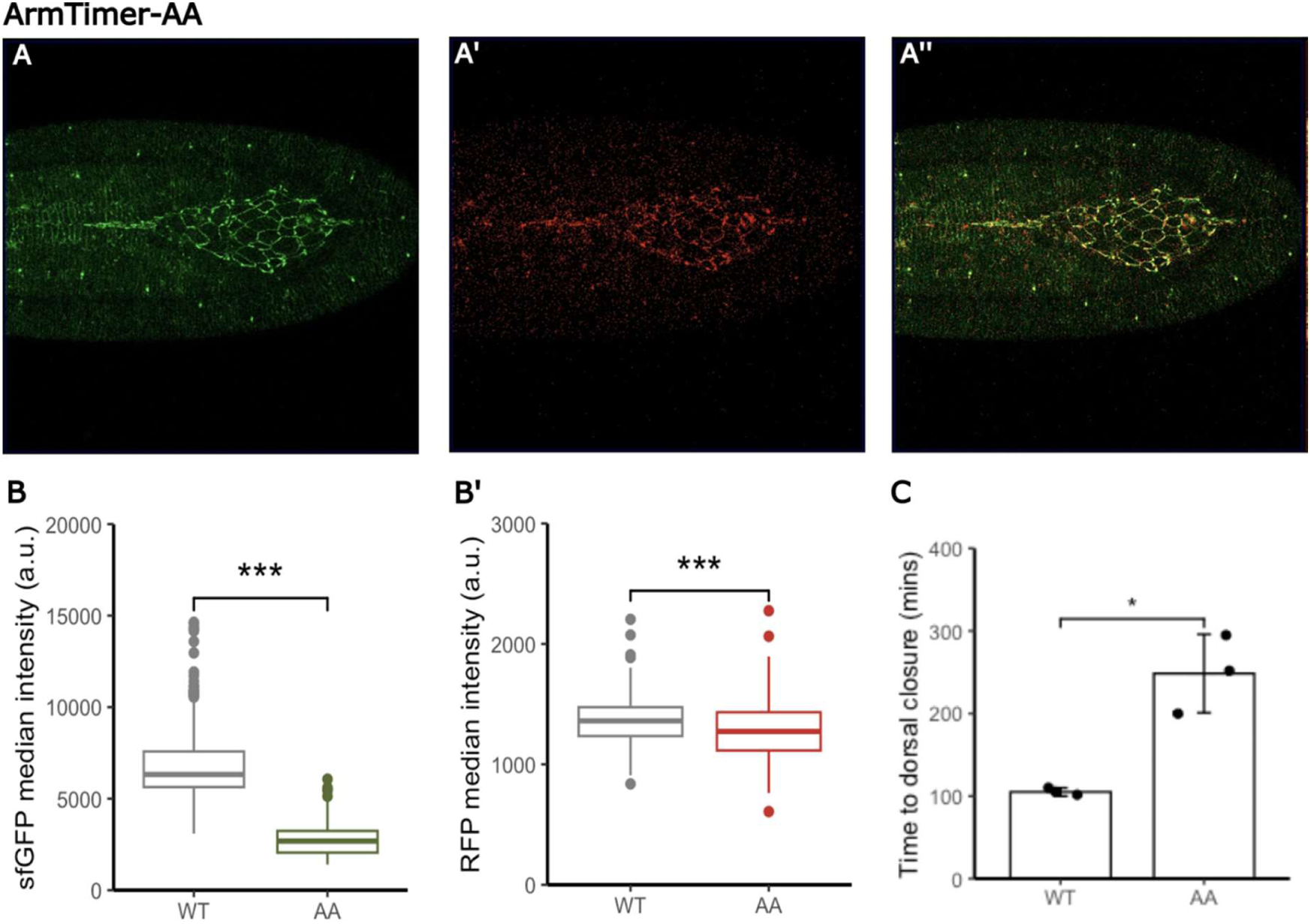
ArmTimer-AA disrupts zippering dynamics, supporting the requirement for Arm–α-catenin interaction. (A–A’’) ArmTimer-AA (T111A, T121A) shows asymmetric zippering and an uneven dorsal gap. (B) GFP comparison indicates lower Arm production/accumulation in ArmTimer-AA relative to wild type at the same time point. (B’) RFP comparison indicates similar stabilization signal to wild type. (C) Dorsal closure duration is significantly increased for ArmTimer-AA.

### Dishevelled regulates Arm during dorsal closure

Dishevelled (Dsh) is a key upstream component of both canonical and non-canonical Wnt signaling pathways that regulate polarity, adhesion, and migration. Null dsh mutants exhibit dorsal closure defects, including dorsal holes.^54^ Canonically, Dsh is recruited to the membrane downstream of Frizzled activation, where it helps recruit Axin and associated kinases to the activating “signalosome,” thereby antagonizing the destruction complex. Non-canonically, Dsh can activate Rho/Rac pathways and downstream JNK signaling, depending on interacting partners and domain usage.^18^

Because we did not have a fluorescently tagged Dsh line suitable for live imaging, we used an indirect “frankenbody” strategy: HA-tagged Dsh is co-expressed with a fluorescent anti-HA nanobody, enabling live visualization of the HA-tagged protein.^67^ Using this approach, we observed that Dsh accumulates in the leading-edge cells where Arm is stabilized (Figure 7A–A’’), placing Dsh in the correct spatial domain to influence Arm dynamics during dorsal closure.

**Figure 7.**
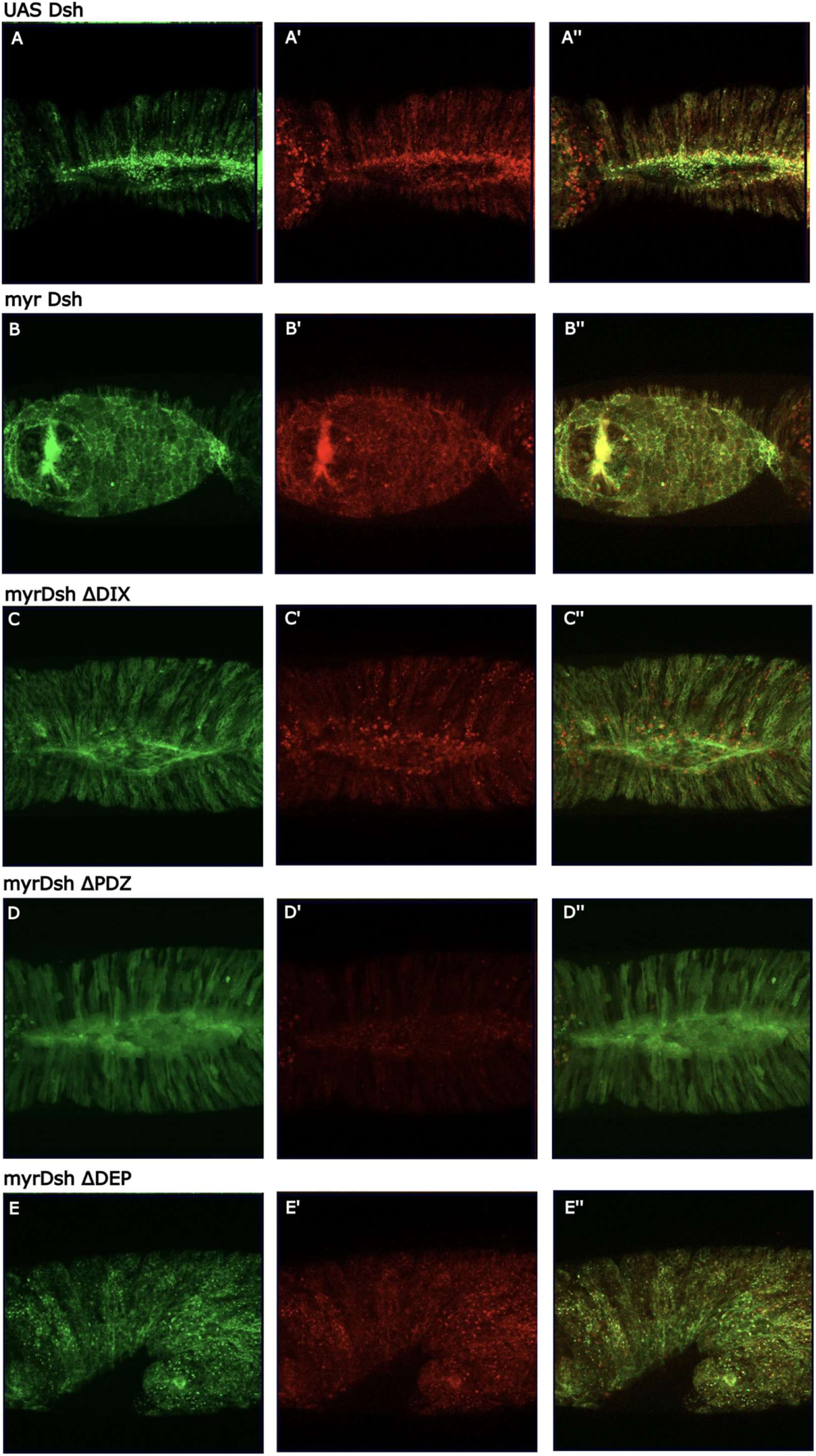
Dishevelled accumulates at the leading edge and the DEP domain is required for dorsal closure. (A–A’’) Dsh visualized with frankenbody reporters accumulates at the leading edge (GFP, RFP, and overlap). (B–B’’) Membrane-tethered Dsh localizes to leading edge and amnioserosa and disrupts amnioserosa junction integrity. (C–C’’) DshΔDIX localizes to membrane and causes amnioserosa integrity defects without blocking closure. (D–D’’) DshΔPDZ is diffuse but dorsal closure proceeds. (E–E’’) DshΔDEP results in failure of germ-band retraction and dorsal closure.

Overexpression of full-length Dsh localized to the leading edge but did not disrupt dorsal closure, consistent with the earlier observation that Wg overexpression does not disrupt closure mechanics. To dissect which Dsh domains are required, we tested membrane-tethered (myristoylated) Dsh constructs with deletions of the conserved DIX, PDZ, and DEP domains.^68^ These domains partition Dsh function across pathway branches: PDZ and DEP support membrane recruitment and Frizzled interaction, whereas DIX supports signalosome assembly and interaction with Axin in canonical signaling. In polarity signaling, DEP is critical for engaging small GTPases and JNK pathway outputs.^69^

- Myr-Dsh localized to the leading edge but disrupted amnioserosa integrity, resembling previously described phenotypes^54^ (Figure 7B).
- Myr-DshΔDIX, which cannot efficiently engage Axin/canonical signalosome formation, localized to membranes and produced amnioserosa integrity defects but did not block dorsal closure (Figure 7C).
- Myr-DshΔPDZ localized more diffusely across the epithelium but dorsal closure proceeded (Figure 7D).
- Myr-DshΔDEP produced severe early defects including defective germ-band retraction and dorsal closure failure (Figure 7E).

Together, these results indicate that the DEP domain is essential for the morphogenetic program culminating in dorsal closure, consistent with a strong role for the Dsh DEP-dependent non-canonical/polarity branch in this process.

### JNK signaling regulates ArmTimer dynamics and is essential for dorsal closure

The requirement for the Dsh DEP domain naturally implicates downstream pathways engaged by DEP, including the Rac/JNK pathway.^69^ The Jun N-terminal kinase (JNK) pathway is a mitogen-activated protein kinase (MAPK) cascade required for dorsal closure, regulating leading-edge behaviors, expression of target genes such as dpp and puc, and actin-based protrusions important for zippering.^70–72^ Genetic studies have shown that JNK pathway mutants fail dorsal closure.^73,74^ However, how JNK activity might translate into Arm stabilization or junctional dynamics remains incompletely defined.

To test whether JNK influences ArmTimer behavior, we modulated JNK levels. Uniform JNK-GFP overexpression did not visibly disrupt dorsal closure, but ArmTimer imaging revealed reduced Arm-RFP intensity, suggesting decreased Arm stabilization or increased turnover at junctions when JNK is elevated (Figure 8A, C). Conversely, JNK knockdown (JNK RNAi) caused a strong early disruption of dorsal closure and altered Arm localization at the leading edge (Figure 8B), consistent with the essential requirement of JNK signaling for this morphogenetic process.

**Figure 8.**
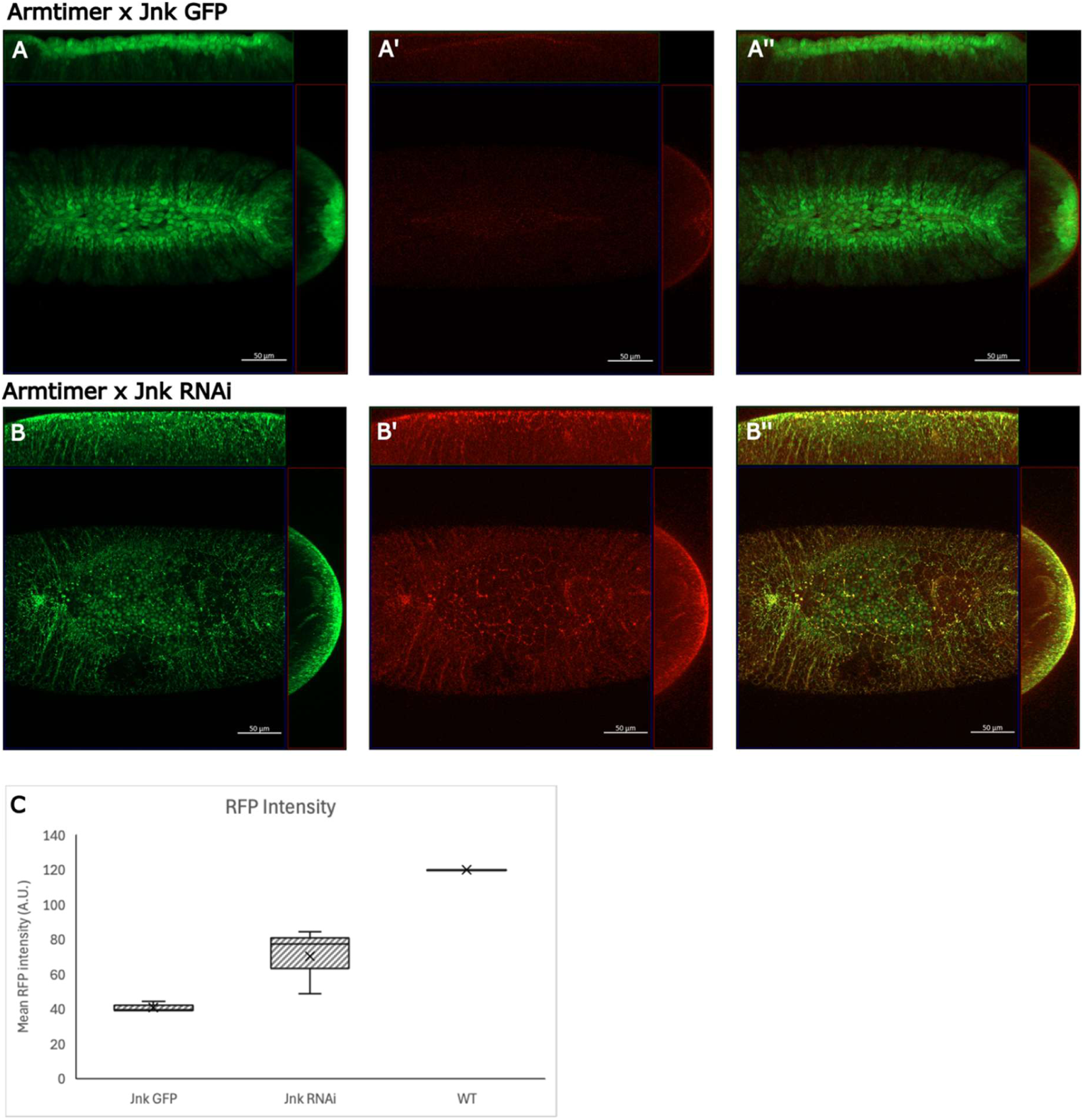
JNK signaling regulates ArmTimer dynamics and is essential for proper dorsal closure. (A–A’’) ArmTimer embryos with uniform JNK-GFP expression show normal closure but reduced Arm-RFP intensity. (B–B’’) ArmTimer × JNK RNAi shows disrupted dorsal closure and altered Arm leading-edge localization. (C) Quantification of Arm-RFP intensity across JNK-GFP, JNK RNAi, and wild-type ArmTimer.

## DISCUSSION

This study uses endogenous protein lifetime reporting and optogenetic perturbation to reveal a striking and unexpected stabilization of β-catenin/Armadillo at adherens junctions during a force-generating morphogenetic process *in vivo*. Given the canonical view that β-catenin stabilization is a hallmark of canonical Wnt pathway activation, we expected Arm stabilization to be most prominent in Wingless-expressing stripes during embryonic patterning. Instead, Arm stabilization is highly enriched at the leading edge during dorsal closure, a mechanically active zone that coordinates epithelial migration, actin cable tension, and zippering.

### Junctional Arm stabilization is essential for dorsal closure mechanics

Our optogenetic inactivation of Arm shows that Arm activity is required for dorsal closure. Importantly, optogenetic inactivation of α-catenin produces a comparable dorsal closure failure phenotype, indicating that adherens junction function is essential for closure. These observations support a model in which dorsal closure requires robust adherens junction coupling to transmit and coordinate forces across cells at the leading edge.

A central mechanistic inference from our mutant analyses is that Arm–α-catenin interaction is critical. Multiple perturbations that compromise this interaction produce dorsal closure defects: α-catenin inactivation (α-cat-CRY2), Arm inactivation (Arm-CRY2), and ΔArm, which lacks the α-catenin binding region, all display failure of zippering forces. ArmTimer-AA, which reduces α-catenin binding through altered threonine residues in the binding region, produces a milder but clear defect—slower and asymmetric zippering. This graded relationship between the extent of interaction disruption and the severity of closure defects strongly supports the conclusion that Arm–α-catenin coupling is required to stabilize junctions and coordinate force transmission during closure.

Threonine phosphorylation of β-catenin/Arm has previously been linked to polarity by stabilizing Arm–α-catenin binding,^66^ and planar polarity in the Drosophila epidermis can be established through junctional Arm dynamics.^16^ Our data extend this concept to the dorsal closure context, showing that compromising α-catenin binding specifically disrupts zippering dynamics, consistent with a requirement for stable, polarized junctions at the leading edge to enable mechanical closure.

### Tyrosine-site mutants suggest altered Arm dynamics without overt failure

Our results with the tyrosine mutants (ArmTimer-EE and ArmTimer-FF) are more ambiguous with respect to a simple, direct model of E-cadherin binding regulation. Both mutants show reduced fluorescence intensity and altered Timer signatures but do not lead to obvious dorsal closure failure. Both tyrosine mutants show reduced stabilization and/or abundance, suggesting these residues influence Arm dynamics. However, the directionality does not map cleanly onto a simple phosphomimetic (less stable) versus phospho-resistant (more stable) expectation.

Prior literature has reported complex and sometimes conflicting roles for tyrosine phosphorylation in regulating β-catenin/E-cadherin affinity and junction stability. Phosphorylation of Y150 reduces affinity for E-cadherin, whereas Y667 has been reported to either increase or decrease affinity depending on context.^49,64,75,76^ In our study, ArmTimer-EE’s preserved RFP but reduced GFP suggests altered production/turnover with a stable fraction maintained, whereas ArmTimer-FF’s reduced RFP suggests reduced stabilization despite normal GFP. These results imply that the tyrosine sites likely influence more than one aspect of Arm behavior— potentially the balance among membrane retention, junction remodeling, and degradation—rather than acting as simple binary switches.

ArmTimer-EE also increases the duration of dorsal closure, suggesting that phosphomimetic Arm sufficiently alters junction dynamics to slow closure without preventing it. Thus, while tyrosine phosphorylation does not appear essential for completion of dorsal closure under our conditions, it likely modulates the kinetics and stability of junctional Arm.

### Dishevelled and JNK provide a link between Wnt components and mechanical morphogenesis

Canonical Wnt activation by Wg overexpression does not perturb dorsal closure or increase leading-edge Arm stability. However, inhibiting Arm stabilization by Axin overexpression causes closure defects, suggesting that the stability machinery regulating Arm abundance remains relevant. At the same time, Dsh accumulates at the leading edge, and perturbing the Dsh DEP domain causes severe morphogenetic failure. Because the DEP domain is central to non-canonical Wnt signaling outputs that engage Rac/JNK pathways, these observations support the notion that Wnt pathway components contribute to dorsal closure primarily through polarity/force-regulatory modules rather than through canonical ligand-driven transcriptional activation.

Consistent with this, JNK knockdown causes severe dorsal closure defects, and JNK overexpression alters ArmTimer RFP levels, suggesting that JNK activity modulates Arm stabilization and/or junction turnover. One interpretation is that increased JNK accelerates junction remodeling and turnover (reducing the stable Arm pool), whereas reduced JNK disrupts cytoskeletal tension, adhesion coordination, and leading-edge dynamics required for closure. Together, our results support a model in which Dsh (via DEP) and JNK signaling regulate junctional Arm stability and/or junction remodeling at the leading edge, thereby providing a functional link between Wnt components and mechanotransduction during dorsal closure.

### Future directions: mechanotransduction and Hippo pathway integration

Since dorsal closure is inherently mechanical, these findings suggest broader links between mechanosensation and mechanotransduction links involving junctions. We observed stable Arm expression in other tissues undergoing mechanotransduction later in embryogenesis, including the gut and trachea (Supplementary Figure S1).^77–79^ This suggests that junctional β-catenin stabilization may be a recurring feature of force-driven morphogenesis. Given that α-catenin has been implicated as a mechanosensor and that the Hippo pathway is force-regulated and central to controlling epithelial growth, a compelling future direction is to test whether Hippo signaling interacts with adherens junction stabilization during dorsal closure. Such research could clarify whether force generation increases Arm stabilization as part of a mechanical reinforcement process or if Arm stabilization itself facilitates effective force transmission and successful closure.

## MATERIALS AND METHODS

### Fly lines, MiMIC/RMCE, and expression of UAS constructs

All crosses for transgene expression were performed as previously described.^31^ Expression of transgenes was driven by the daughterless, C381, and armadillo-GAL4 drivers (Kaur et al., 2017b). Additional stocks were obtained from the Bloomington Drosophila Stock Center (NIH P40OD018537). The tandem fluorescent protein timer recombination insertion construct was synthesized by SynbioTech (Monmouth Junction, NJ, USA) and injected by Bestgene (Chino Hills, CA, USA) following MiMIC-RMCE protocols.^27–29,80^ and timer protocols.^40,81^

| Fly Line | Reference |
| --- | --- |
| ArmTimer: Mi[MIC]armMI08675 - RMCE sfGFP-RFP Phase 1 | This study |
| OptoArm: armMI08675 MiMIC RMCE Cry2-mCherry Phase 1 | This study |
| Mi(PT-GFSTF.1)armMI08675-GFSTF | BDSC 60561 |
| ArmTimerWT: Mi[MIC]armMI08675 - RMCE ArmWT - sfGFP-RFP | This study |
| ArmTimerFF: Mi[MIC]armMI08675 - RMCE ArmY150F, Y667F - sfGFP-RFP | This study |
| ArmTimerEE: Mi[MIC]armMI08675 - RMCE ArmY150E, Y667E - sfGFP-RFP | This study |
| ArmTimerAA: Mi[MIC]armMI08675 - RMCE ArmT111A, T121A - sfGFP-RFP | This study |
| ArmTimerF1a: Mi[MIC]armMI08675 - RMCE ArmR394H - sfGFP-RFP | This study |
| OptoαCat: Mi[MIC]α-CatMI02577 - RMCE Cry2-mCherry Phase 0 | This study |
| OptoDlg: Mi[MIC]dlg1MI06353 - RMCE Cry2-mCherry Phase 0 | This study |
| UAS- LifeActin-mKO | This study |
| Arm RMCE CRY2-mCh, UbiGFP, FRT19A (for germline clones) | This study |
| UAS Franken (anti-HA epitope nanobody)-GFP-mCherry | Murakawa et al, 2022<sup>82</sup> |
| UAS myr-Dsh-HA | Kaur et al, 2017<sup>68</sup> |
| UAS myr-Dsh ΔDIX-HA | Kaur et al, 2017 |
| UAS myr-Dsh ΔPDZ-HA | Kaur et al, 2017 |
| UAS myr-Dsh ΔDEP-HA | Kaur et al, 2017 |
| UAS-Dsh-HA | Kaur et al, 2017 |
| UAS-Wingless | BDSC 2075 |
| UAS-RFP | BDSC 30555 |
| UAS-Axin-Cry2-mCherry | Bunnag et al. 2020<sup>35</sup> |
| UAS-ΔArm | Zecca et al. 1996<sup>60</sup> |
| UAS-ArmS56A-sfGFP-RFP | This study |
| UAS-E-cadherin RFP | Gift Timothy Saunders |

The frankenbody approach was used to visualize overexpression and mutants of Dishevelled.^67^ This system uses an α-HA nanobody fused to GFP, mCherry, or both, co-expressed with any HA-tagged transgene, enabling live imaging of HA-tagged proteins.^82^ Transgene expression was driven using the daughterless (da), C381, and armadillo (arm)-GAL4 drivers. Unless otherwise indicated, all additional fly stocks were obtained from the Bloomington Drosophila Stock Center (NIH P40OD018537).

The tandem fluorescent protein timer recombination insertion construct was synthesized by SynbioTech (Monmouth Junction, NJ, USA) and injected by Bestgene (Chino Hills, CA, USA).

### Imaging

Arm-Timer embryos were imaged using a Zeiss Lightsheet Z.1 microscope (Carl Zeiss, Germany). Embryos were dechorionated with bleach, rinsed twice in water, dried, and loaded into a capillary containing 1% low-melting agarose (Type VII-A) in water (Sigma-Aldrich, St. Louis, MO, USA).^31,83^ For imaging, the agarose plug was extruded from the capillary, and the sample was suspended in the imaging chamber filled with water. Images were acquired using a water-immersion W Plan-Apochromat 40×/1.0 UV-VIS detection objective (Carl Zeiss, Jena, Germany). Samples were imaged every 5 minutes for 200 cycles, using 5% laser power at 488 nm and 15% laser power at 561 nm. A late developmental time point was selected for analysis. Background fluorescence was automatically calculated in ZEN 2014 using the background correction function (Carl Zeiss). Deep, out-of-focus z-stacks were excluded, and maximum intensity projections were generated from in-focus stacks for each embryo. Embryos were dechorionated with bleach, rinsed twice with water, and mounted in water in an imaging dish.

Dorsal closure imaging was performed on a Zeiss LSM 800 (Carl Zeiss, Germany) with acquisition every 5 min for 60 cycles, using 3% laser power at 488 nm and 5% laser power at 561 nm. Dishevelled imaging was performed on the same system using 5% laser power at 488 nm and 10% laser power at 561 nm, also every 5 min for 60 cycles. Images were processed using the maximum intensity projection function in ZEN 2014 SP (Carl Zeiss). RFP and GFP cluster intensities were quantified using IMARIS 9.0 (Bitplane AG, UK). Z-stacks from the LSM800 were rendered using the Surface module, and fluorescence intensities were extracted for plotting. All time points were analyzed, and all channels were rendered using identical brightness and sensitivity settings. Statistical analyses were performed in R (v4.2.2), and plots were generated using ggpubr (v0.6.0).

## Author Contributions

Conceptualization, N.S.T.; methodology, A.O., S.S., P.K., P.T., N.S.T; investigation, A.O., S.S., P.K., P.T.; writing—original draft preparation, A.O., and N.S.T.; writing—review and editing A.O., S.S., P.K., P.T., N.S.T,; supervision and funding acquisition, N.S.T. All authors have read and agreed to the published version of the manuscript.

## Funding

This work was supported by the Singapore Ministry of Education, AcRF grants IG21-SG103 and FY2022-MOET1-0003 to N.S.T.

## Declaration of generative AI and AI-assisted technologies in the writing process

During the preparation of this work, the authors used OpenAI GPT-5.2 to improve the manuscript’s readability. After using this tool/service, the authors reviewed and edited the content as needed and take full responsibility for the content of the published article.

## Acknowledgments

Stocks obtained from the Bloomington Drosophila Stock Center (NIH P40OD018537) were used in this study. We used Flybase.org release FB2026_01.^84^

## Conflicts of Interest

The authors declare no conflicts of interest.

## Notes

### Competing Interest Statement

The authors have declared no competing interest.

## References

1. Kemler, R., Babinet, C., Eisen, H., and Jacob, F. (1977). Surface antigen in early differentiation. Proceedings of the National Academy of Sciences 74, 4449–4452.

2. Takeichi, M. (1977). Functional correlation between cell adhesive properties and some cell surface proteins. The Journal of cell biology 75, 464–474.

3. Nagafuchi, A., and Takeichi, M. (1988). Cell binding function of E-cadherin is regulated by the cytoplasmic domain. The EMBO journal 7, 3679–3684.

4. Ozawa, M., Baribault, H., and Kemler, R. (1989). The cytoplasmic domain of the cell adhesion molecule uvomorulin associates with three independent proteins structurally related in different species. The EMBO journal 8, 1711–1717.

5. Herrenknecht, K., Ozawa, M., Eckerskorn, C., Lottspeich, F., Lenter, M., and Kemler, R. (1991). The uvomorulin-anchorage protein alpha catenin is a vinculin homologue. Proceedings of the National Academy of Sciences 88, 9156–9160.

6. Nagafuchi, A., Takeichi, M., and Tsukita, S. (1991). The 102 kd cadherin-associated protein: similarity to vinculin and posttranscriptional regulation of expression. Cell 65, 849–857.

7. Nagafuchi, A., Ishihara, S., and Tsukita, S. (1994). The roles of catenins in the cadherin-mediated cell adhesion: functional analysis of E-cadherin-alpha catenin fusion molecules. J Cell Biol 127, 235–245. 10.1083/jcb.127.1.235.

8. Peifer, M., Pai, L.M., and Casey, M. (1994). Phosphorylation of the Drosophila adherens junction protein Armadillo: roles for wingless signal and zeste-white 3 kinase. Dev Biol 166, 543–556. 10.1006/dbio.1994.1336.

9. Wieschaus, E., Nüsslein-Volhard, C., and Jürgens, G. (1984). Mutations affecting the pattern of the larval cuticle inDrosophila melanogaster. Wilhelm Roux’s archives of developmental biology 193, 296–307. 10.1007/BF00848158.

10. Riggleman, B., Schedl, P., and Wieschaus, E. (1990). Spatial expression of the Drosophila segment polarity gene armadillo is posttranscriptionally regulated by wingless. Cell 63, 549–560.

11. Wieschaus, E., and Riggleman, R. (1987). Autonomous requirements for the segment polarity gene armadillo during Drosophila embryogenesis. Cell 49, 177–184.

12. Nüsslein-Volhard, C., and Wieschaus, E. (1980). Mutations affecting segment number and polarity in Drosophila. Nature 287, 795–801.

13. Wieschaus, E., and Nüsslein-Volhard, C. (2016). The Heidelberg screen for pattern mutants of Drosophila: a personal account. Annual review of cell and developmental biology 32, 1–46.

14. Zallen, J.A., and Wieschaus, E. (2004). Patterned gene expression directs bipolar planar polarity in Drosophila. Developmental cell 6, 343–355.

15. Blankenship, J.T., Backovic, S.T., Sanny, J.S., Weitz, O., and Zallen, J.A. (2006). Multicellular rosette formation links planar cell polarity to tissue morphogenesis. Developmental cell 11, 459–470.

16. Colosimo, P.F., and Tolwinski, N.S. (2006). Wnt, Hedgehog and junctional Armadillo/beta-catenin establish planar polarity in the Drosophila embryo. PLoS One 1, e9. 10.1371/journal.pone.0000009.

17. Kaplan, N.A., and Tolwinski, N.S. (2010). Spatially defined Dsh-Lgl interaction contributes to directional tissue morphogenesis. J Cell Sci 123, 3157–3165. 10.1242/jcs.069898.

18. Schlessinger, K., Hall, A., and Tolwinski, N. (2009). Wnt signaling pathways meet Rho GTPases. Genes Dev 23, 265–277. 10.1101/gad.1760809.

19. Liu, S., Haghani, S., Petretto, E., Madan, B., Harmston, N., and Virshup, D.M. Identification of Wnt-regulated genes that are repressed by, or independent of, β-catenin. The FEBS Journal n/a. 10.1111/febs.70417.

20. Bertet, C., Sulak, L., and Lecuit, T. (2004). Myosin-dependent junction remodelling controls planar cell intercalation and axis elongation. Nature 429, 667–671.

21. Drees, F., Pokutta, S., Yamada, S., Nelson, W.J., and Weis, W.I. (2005). α-catenin is a molecular switch that binds E-cadherin-β-catenin and regulates actin-filament assembly. Cell 123, 903–915.

22. Etienne-Manneville, S., and Hall, A. (2003). Cdc42 regulates GSK-3β and adenomatous polyposis coli to control cell polarity. Nature 421, 753–756.

23. Knoblich, J.A. (2008). Mechanisms of asymmetric stem cell division. Cell 132, 583–597.

24. Nelson, W.J., and Weis, W.I. (2016). 25 years of tension over actin binding to the cadherin cell adhesion complex: the devil is in the details. Trends in cell biology 26, 471–473.

25. Yamada, S., Pokutta, S., Drees, F., Weis, W.I., and Nelson, W.J. (2005). Deconstructing the cadherin-catenin-actin complex. Cell 123, 889–901.

26. Kanca, O., Bellen, H.J., and Schnorrer, F. (2017). Gene tagging strategies to assess protein expression, localization, and function in Drosophila. Genetics 207, 389–412.

27. Li-Kroeger, D., Kanca, O., Lee, P.T., Cowan, S., Lee, M.T., Jaiswal, M., Salazar, J.L., He, Y., Zuo, Z., and Bellen, H.J. (2018). An expanded toolkit for gene tagging based on MiMIC and scarless CRISPR tagging in Drosophila. Elife 7, e38709. 10.7554/eLife.38709.

28. Nagarkar-Jaiswal, S., DeLuca, S.Z., Lee, P.-T., Lin, W.-W., Pan, H., Zuo, Z., Lv, J., Spradling, A.C., and Bellen, H.J. (2015). A genetic toolkit for tagging intronic MiMIC containing genes. Elife 4, e08469.

29. Nagarkar-Jaiswal, S., Lee, P.-T., Campbell, M.E., Chen, K., Anguiano-Zarate, S., Cantu Gutierrez, M., Busby, T., Lin, W.-W., He, Y., and Schulze, K.L. (2015). A library of MiMICs allows tagging of genes and reversible, spatial and temporal knockdown of proteins in Drosophila. elife *4*, e05338.

30. Venken, K.J., Schulze, K.L., Haelterman, N.A., Pan, H., He, Y., Evans-Holm, M., Carlson, J.W., Levis, R.W., Spradling, A.C., Hoskins, R.A., and Bellen, H.J. (2011). MiMIC: a highly versatile transposon insertion resource for engineering Drosophila melanogaster genes. Nat Methods 8, 737–743. 10.1038/nmeth.1662.

31. Kaur, P., Saunders, T.E., and Tolwinski, N.S. (2017). Coupling optogenetics and light-sheet microscopy, a method to study Wnt signaling during embryogenesis. Sci Rep 7, 16636. 10.1038/s41598-017-16879-0.

32. Lim, C.H., Kaur, P., Teo, E., Lam, V.Y.M., Zhu, F., Kibat, C., Gruber, J., Mathuru, A.S., and Tolwinski, N.S. (2020). Application of optogenetic Amyloid-beta distinguishes between metabolic and physical damages in neurodegeneration. Elife 9. 10.7554/eLife.52589.

33. Yadav, V., Tolwinski, N., and Saunders, T.E. (2021). Spatiotemporal sensitivity of mesoderm specification to FGFR signalling in the Drosophila embryo. Sci Rep 11, 14091. 10.1038/s41598-021-93512-1.

34. Suresh, J., Khor, I.W., Kaur, P., Heng, H.L., Torta, F., Dawe, G.S., Tai, E.S., and Tolwinski, N.S. (2020). Shared signaling pathways in Alzheimer’s and metabolic disease may point to new treatment approaches. FEBS J. 10.1111/febs.15540.

35. Bunnag, N., Tan, Q.H., Kaur, P., Ramamoorthy, A., Sung, I.C.H., Lusk, J., and Tolwinski, N.S. (2020). An Optogenetic Method to Study Signal Transduction in Intestinal Stem Cell Homeostasis. J Mol Biol 432, 3159–3176. 10.1016/j.jmb.2020.03.019.

36. Ambrosi, G., Voloshanenko, O., Eckert, A.F., Kranz, D., Nienhaus, G.U., and Boutros, M. (2022). Allele-specific endogenous tagging and quantitative analysis of β-catenin in colorectal cancer cells. Elife 11. 10.7554/eLife.64498.

37. de Man, S.M., Zwanenburg, G., van der Wal, T., Hink, M.A., and van Amerongen, R. (2021). Quantitative live-cell imaging and computational modeling shed new light on endogenous WNT/CTNNB1 signaling dynamics. Elife 10. 10.7554/eLife.66440.

38. Barry, J.D., Dona, E., Gilmour, D., and Huber, W. (2016). TimerQuant: a modelling approach to tandem fluorescent timer design and data interpretation for measuring protein turnover in embryos. Development 143, 174–179. 10.1242/dev.125971.

39. Durrieu, L., Kirrmaier, D., Schneidt, T., Kats, I., Raghavan, S., Hufnagel, L., Saunders, T.E., and Knop, M. (2018). Bicoid gradient formation mechanism and dynamics revealed by protein lifetime analysis. Molecular Systems Biology 14, e8355. 10.15252/msb.20188355.

40. Khmelinskii, A., Keller, P.J., Bartosik, A., Meurer, M., Barry, J.D., Mardin, B.R., Kaufmann, A., Trautmann, S., Wachsmuth, M., Pereira, G., et al. (2012). Tandem fluorescent protein timers for in vivo analysis of protein dynamics. Nat Biotechnol 30, 708–714. 10.1038/nbt.2281.

41. Revenu, C., Streichan, S., Dona, E., Lecaudey, V., Hufnagel, L., and Gilmour, D. (2014). Quantitative cell polarity imaging defines leader-to-follower transitions during collective migration and the key role of microtubule-dependent adherens junction formation. Development 141, 1282–1291. 10.1242/dev.101675.

42. Tan, Q.H., Otgonbaatar, A., Kaur, P., Ga, A.F., Harmston, N.P., and Tolwinski, N.S. (2024). The Wnt Co-Receptor PTK7/Otk and Its Homolog Otk-2 in Neurogenesis and Patterning. Cells 13. 10.3390/cells13050365.

43. Cadigan, K.M., and Nusse, R. (1997). Wnt signaling: a common theme in animal development. Genes Dev 11, 3286–3305. 10.1101/gad.11.24.3286.

44. Lee, E., Salic, A., Krüger, R., Heinrich, R., and Kirschner, M.W. (2003). The roles of APC and Axin derived from experimental and theoretical analysis of the Wnt pathway. PLoS Biol 1, E10. 10.1371/journal.pbio.0000010.

45. Moldaver, S., Thibeault, P.E., Robitaille, M., Au, A., Lin, S., MacLeod, G., Junge, H.J., Gammons, M.V., Yip, C.M., and Angers, S. (2026). Wnt-dependent Frizzled clustering is required for Dishevelled phosphorylation but insufficient for β-catenin stabilization. Science Signaling 19, eaec0204. doi:10.1126/scisignal.aec0204.

46. Gammons, M., and Bienz, M. (2018). Multiprotein complexes governing Wnt signal transduction. Current Opinion in Cell Biology 51, 42–49. 10.1016/j.ceb.2017.10.008.

47. Maurice, M.M., and Angers, S. (2025). Mechanistic insights into Wnt–β-catenin pathway activation and signal transduction. Nature Reviews Molecular Cell Biology 26, 371–388. 10.1038/s41580-024-00823-y.

48. Lee, M.H., Koria, P., Qu, J., and Andreadis, S.T. (2009). JNK phosphorylates beta-catenin and regulates adherens junctions. Faseb j 23, 3874–3883. 10.1096/fj.08-117804.

49. Roura, S., Miravet, S., Piedra, J., de Herreros, A.G., and Dunach, M. (1999). Regulation of E-cadherin/Catenin association by tyrosine phosphorylation. Journal of Biological Chemistry 274, 36734–36740.

50. Hayes, P., and Solon, J. (2017). Drosophila dorsal closure: An orchestra of forces to zip shut the embryo. Mech Dev 144, 2–10. 10.1016/j.mod.2016.12.005.

51. Kiehart, D.P., Crawford, J.M., Aristotelous, A., Venakides, S., and Edwards, G.S. (2017). Cell sheet morphogenesis: dorsal closure in Drosophila melanogaster as a model system. Annual review of cell and developmental biology 33, 169–202.

52. Kiehart, D.P., Galbraith, C.G., Edwards, K.A., Rickoll, W.L., and Montague, R.A. (2000). Multiple Forces Contribute to Cell Sheet Morphogenesis for Dorsal Closure in Drosophila. Journal of Cell Biology 149, 471–490. 10.1083/jcb.149.2.471.

53. Gorfinkiel, N., and Arias, A.M. (2007). Requirements for adherens junction components in the interaction between epithelial tissues during dorsal closure in Drosophila. Journal of cell science 120, 3289–3298.

54. Morel, V., and Arias, A.M. (2004). Armadillo/β-catenin-dependent Wnt signalling is required for the polarisation of epidermal cells during dorsal closure in Drosophila. Development 131, 3273–3283.

55. McEwen, D.G., Cox, R.T., and Peifer, M. (2000). The canonical Wg and JNK signaling cascades collaborate to promote both dorsal closure and ventral patterning. Development 127, 3607–3617.

56. Tolwinski, N.S., and Wieschaus, E. (2004). A nuclear function for armadillo/beta-catenin. PLoS Biol 2, E95. 10.1371/journal.pbio.0020095.

57. Tolwinski, N.S., and Wieschaus, E. (2001). Armadillo nuclear import is regulated by cytoplasmic anchor Axin and nuclear anchor dTCF/Pan. Development 128, 2107–2117. 10.1242/dev.128.11.2107.

58. Rijsewijk, F., Schuermann, M., Wagenaar, E., Parren, P., Weigel, D., and Nusse, R. (1987). The Drosophila homology of the mouse mammary oncogene <em>int</em>-1 is identical to the segment polarity gene <em>wingless</em>. Cell 50, 649–657. 10.1016/0092-8674(87)90038-9.

59. Tolwinski, N.S., Wehrli, M., Rives, A., Erdeniz, N., DiNardo, S., and Wieschaus, E. (2003). Wg/Wnt signal can be transmitted through arrow/LRP5,6 and Axin independently of Zw3/Gsk3beta activity. Dev Cell 4, 407–418. 10.1016/s1534-5807(03)00063-7.

60. Zecca, M., Basler, K., and Struhl, G. (1996). Direct and Long-Range Action of a Wingless Morphogen Gradient. Cell 87, 833–844. 10.1016/S0092-8674(00)81991-1.

61. Piedra, J., Martinez, D., Castano, J., Miravet, S., Dunach, M., and de Herreros, A.G. (2001). Regulation of beta-catenin structure and activity by tyrosine phosphorylation. J Biol Chem 276, 20436–20443. 10.1074/jbc.M100194200.

62. Piedra, J., Martínez, D., Castaño, J., Miravet, S., Duñach, M., and García de Herreros, A. (2016). Regulation of β-catenin structure and activity by tyrosine phosphorylation. J Biol Chem 291, 11463. 10.1074/jbc.A116.100194.

63. Tamada, M., Farrell, D.L., and Zallen, J.A. (2012). Abl regulates planar polarized junctional dynamics through β-catenin tyrosine phosphorylation. Developmental cell 22, 309–319.

64. Tominaga, J., Fukunaga, Y., Abelardo, E., and Nagafuchi, A. (2008). Defining the function of β-catenin tyrosine phosphorylation in cadherin-mediated cell–cell adhesion. Genes to Cells 13, 67–77. 10.1111/j.1365-2443.2007.01149.x.

65. van Veelen, W., Le, N.H., Helvensteijn, W., Blonden, L., Theeuwes, M., Bakker, E.R., Franken, P.F., van Gurp, L., Meijlink, F., van der Valk, M.A., et al. (2011). beta-catenin tyrosine 654 phosphorylation increases Wnt signalling and intestinal tumorigenesis. Gut 60, 1204–1212. 10.1136/gut.2010.233460.

66. Bek, S., and Kemler, R. (2002). Protein kinase CKII regulates the interaction of beta-catenin with alpha-catenin and its protein stability. J Cell Sci 115, 4743–4753. 10.1242/jcs.00154.

67. Zhao, N., Kamijo, K., Fox, P.D., Oda, H., Morisaki, T., Sato, Y., Kimura, H., and Stasevich, T.J. (2019). A genetically encoded probe for imaging nascent and mature HA-tagged proteins in vivo. Nat Commun 10, 2947. 10.1038/s41467-019-10846-1.

68. Kaur, P., Lam, V.Y.M., Mannava, A.G., Suresh, J., Jenny, A., and Tolwinski, N.S. (2017). Membrane Targeting of Disheveled Can Bypass the Need for Arrow/LRP5. Sci Rep 7, 6934. 10.1038/s41598-017-04414-0.

69. Boutros, M., Paricio, N., Strutt, D.I., and Mlodzik, M. (1998). Dishevelled activates JNK and discriminates between JNK pathways in planar polarity and wingless signaling. Cell 94, 109–118. 10.1016/s0092-8674(00)81226-x.

70. Harden, N. (2002). Signaling pathways directing the movement and fusion of epithelial sheets: lessons from dorsal closure in Drosophila. Differentiation: REVIEW 70, 181–203.

71. Noselli, S., and Agnès, F. (1999). Roles of the JNK signaling pathway in Drosophila morphogenesis. Current opinion in genetics & development 9, 466–472.

72. Jacinto, A., Wood, W., Balayo, T., Turmaine, M., Martinez-Arias, A., and Martin, P. (2000). Dynamic actin-based epithelial adhesion and cell matching during Drosophila dorsal closure. Curr Biol 10, 1420–1426. 10.1016/s0960-9822(00)00796-x.

73. Riesgo-Escovar, J.R., Jenni, M., Fritz, A., and Hafen, E. (1996). The Drosophila Jun-N-terminal kinase is required for cell morphogenesis but not for DJun-dependent cell fate specification in the eye. Genes Dev 10, 2759–2768. 10.1101/gad.10.21.2759.

74. Homsy, J.G., Jasper, H., Peralta, X.G., Wu, H., Kiehart, D.P., and Bohmann, D. (2006). JNK signaling coordinates integrin and actin functions during Drosophila embryogenesis. Dev Dyn 235, 427–434. 10.1002/dvdy.20649.

75. Hamada-Kawaguchi, N., Nishida, Y., and Yamamoto, D. (2015). Btk29A-mediated tyrosine phosphorylation of armadillo/β-catenin promotes ring canal growth in Drosophila oogenesis. PLoS One 10, e0121484. 10.1371/journal.pone.0121484.

76. Shomori, K., Ochiai, A., Akimoto, S., Ino, Y., Shudo, K., Ito, H., and Hirohashi, S. (2009). Tyrosine-phosphorylation of the 12th armadillo-repeat of β-catenin is associated with cadherin dysfunction in human cancer. International journal of oncology 35, 517–524.

77. Caussinus, E., Colombelli, J., and Affolter, M. (2008). Tip-Cell Migration Controls Stalk-Cell Intercalation during <em>Drosophila</em> Tracheal Tube Elongation. Current Biology 18, 1727–1734. 10.1016/j.cub.2008.10.062.

78. Farge, E. (2003). Mechanical induction of Twist in the Drosophila foregut/stomodeal primordium. Current biology 13, 1365–1377.

79. Houtekamer, R.M., van der Net, M.C., Maurice, M.M., and Gloerich, M. (2022). Mechanical forces directing intestinal form and function. Current Biology 32, R791–R805.

80. Lee, P.-T., Zirin, J., Kanca, O., Lin, W.-W., Schulze, K.L., Li-Kroeger, D., Tao, R., Devereaux, C., Hu, Y., and Chung, V. (2018). A gene-specific T2A-GAL4 library for Drosophila. elife 7, e35574.

81. Khmelinskii, A., and Knop, M. (2014). Analysis of protein dynamics with tandem fluorescent protein timers. In Exocytosis and Endocytosis, (Springer), pp. 195–210.

82. Murakawa, T., Nakamura, T., Kawaguchi, K., Murayama, F., Zhao, N., Stasevich, T.J., Kimura, H., and Fujita, N. (2022). A Drosophila toolkit for HA-tagged proteins unveils a block in autophagy flux in the last instar larval fat body. Development 149. 10.1242/dev.200243.

83. Kaur, P., Kibat, C., Teo, E., Gruber, J., Mathuru, A., and Tolwinski, A.N.S. (2020). Use of Optogenetic Amyloid-beta to Monitor Protein Aggregation in Drosophila melanogaster, Danio rerio and Caenorhabditis elegans. Bio Protoc 10, e3856. 10.21769/BioProtoc.3856.

84. Öztürk-Çolak, A., Marygold, S.J., Antonazzo, G., Attrill, H., Goutte-Gattat, D., Jenkins, V.K., Matthews, B.B., Millburn, G., dos Santos, G., Tabone, C.J., and Consortium, F. (2024). FlyBase: updates to the Drosophila genes and genomes database. Genetics 227. 10.1093/genetics/iyad211.

